# Agent SPI-WSI: In context learning for computationally spatial pathway inferring on whole slide histopathology images conditioned on bulk RNA sequencing using pathologist in the loop

**DOI:** 10.1101/2025.10.16.682972

**Authors:** Rajat Vashistha, Sandra Brosda, Clemence J. Belle, Lauren G. Aoude, Nic Waddell, Soumen Ghosh, Caroline Cooper, Andrew P. Barbour, Viktor Vegh

## Abstract

Bulk RNA sequencing, while cost-effective compared to high-resolution spatial transcriptomics, averages gene expression across heterogeneous cell populations and thus lacks spatial context. To address this limitation, we introduce a structured, human–guided, multi-stage computational AI agent SPI-WSI, that iteratively generates, evaluates, and refines biologically informative natural-language prompts, thereby localizing bulk-derived pathway activity within histopathology slides. Our pipeline uses in-context prompting in large language models (LLMs) to adapt dynamically to the task of prompt generation. Candidate prompts are first produced by the LLM and then subjected to a secondary pathologist critiquing by the same LLM that cross-references PubMed to ensure both biological plausibility and specificity. Each approved prompt against their image tile is scored using the vision language foundation model (CONCH). We benchmarked different LLMs, Gemini 2.0, Gemini 2.5, Claude 3.7 and Claude 4.0, and found that Claude 4.0 achieves the highest cosine similarity (0.7) between image and prompt embeddings. Pathologist-driven scoring and manual segmentation confirm that our method accurately identifies clusters of spatial pathological morphologies. In addition, we have validated the method against ground-truth spatial transcriptomic spots using inhouse and public datasets. Overall, the trend emphasized by ground-truth spatial RNA sequencing prompts is closely aligned with those from bulk prompt. This pathologist-in-the-loop workflow enables large-scale, reproducible tissue profiling and grounds AI-driven spatial annotations.

## 1. Introduction

Spatial transcriptomics (ST) provides spatially resolved quantification of gene expression within tissue sections, integrating molecular measurements with histological context at near-cellular resolution [21]. This coupling is essential for linking molecular signatures to tissue architecture and for interrogating tumour heterogeneity, cell–cell interactions, and spatially organised signalling programmes [5]. Recent developments at the interface of ST and digital histopathology have substantially advanced our understanding of tissue structure, cellular crosstalk, and molecular diversity across multiple cancers [9].

Notwithstanding these advantages, wider adoption of ST in research and clinical workflows remains limited by high costs, technical complexity, and labour-intensive protocols [15]. Machine learning particularly deep learning is increasingly incorporated into spatial analyses and shows promise for enhancing interpretation [8, 9]. Several studies now infer gene expression directly from histopathology, yielding insights into tumour heterogeneity and potential therapeutic targets [14, 19]. Related weakly supervised approaches infer molecular characteristics such as DNA mutations, RNA expression, and DNA methylation from wholeslide images; however, fine-grained spatial profiling is still nascent owing to data scarcity and methodological constraints.

Oesophageal cancer is a highly heterogeneous malignancy [1] with poor outcomes and limited therapeutic options [2, 4, 11]. Actionable biomarkers remain scarce, and immunotherapy has conferred benefit only to small patient subsets in recent trials [7, 16]. Resolving the spatial landscape of these tumours is therefore critical for biomarker discovery and patient stratification.

Early computational efforts combine spot-level tissue images with deep architectures to predict local gene expression [20]. For example, M2ORT, a transformer-based regression model, captures hierarchical tissue features and reports strong performance [17]. PH2ST reduces dependence on large paired WSI–ST datasets by leveraging a limited number of annotated spots and cross-attention for ST prompt-guided representation refinement [13], although paired measurements remain necessary. GHIST extends prediction towards subcellular resolution by training on emerging single-cell ST technologies [3].

Large language models (LLMs) further expand the methodological landscape by enabling reasoning, planning, and tool integration [18]. SpatialAgent illustrates an LLM-based system that autonomously orchestrates spatial-biology workflows from experimental design to multimodal analysis and hypothesis generation—via adaptive reasoning and dynamic tool use rather than fixed pipelines [18]. Nevertheless, it does not directly align transcriptomic signals with histology. LLM’s use of in-context prompting has emerged as a powerful paradigm for dynamic task adaptation without parameter updates [12].

By conditioning model behaviour on domain-specific instructions embedded in the input prompt, in-context learning can produce contextually precise, semantically aligned outputs. This is particularly valuable in biomedical imaging, where integrating structured knowledge with domain-specific terminology is essential for interpretability and clinical relevance.

Unlike prior spatial-transcriptomics predictors (e.g., TRIPLEX, PH2ST, GHIST, Stem), which are data dependent and train fixed architectures to regress spot-or cell-level expression, our framework reformulates the problem as an AI–agent–driven, patient-specific prompt generation and pathway localization task. Rather than directly competing on per-gene regression, it provides a cost-effective, interpretable layer that operates with only bulk RNA-seq and the WSI, and can be used stand-alone to triage histopathology slides or alongside regression models to guide, constrain, or audit their spatial outputs. The proposed methodology integrates LLM, a pretrained visual encoder specifically trained on histopathological image tiles [10], and a human-in-the-loop feedback mechanism, effectively simulating expert-driven iterative refinement.

Specifically, we hypothesised that if normalised pathway activities can induce characteristic histopathologic patterns and uses prompt engineering to deconvolute the bulk RNA signal into these spatially localised patterns. Code will be open sourced upon publication. Our main contributions are as follows:

- We propose a framework for AI agent based patient-specific prompt generation that aligns transcriptomic pathway activity with spatially localized histology features.
- We integrated a customizable penalization strategy and a human-in-the-loop feedback mechanism with structured conversational memory to iteratively refine generative language model outputs.
- We utilized dynamic PubMed-informed critique and semantic categorization to evaluate, rewrite, and score generated prompts based on biomedical evidence.
- We compared the performance of the state of the art LLMs Gemini and Claude and validated the proposed model using spatial data.

## 2. Methods

### 2.1. Dataset and Preprocessing

Oesophageal cancer tissue samples were collected from 130 patients with written informed consent. These samples were used for both WSI and bulk RNA sequencing analysis. Samples were collected at endoscopy and stored in RNAlater. RNA was extracted using the Qiagen AllPrep DNA/RNA mini kit according to standard protocol (Qiagen, Germany) and sequenced on a NovaseqX (150 bp paired-end; Illumina) after library preparation (TruSeq Stranded mRNA Library Prep Kit). Sequencing data was aligned to human reference genome GRCh38 using STAR (v2.5.2a). Sequencing adaptors were trimmed with Cutadapt (v1.9) and gene annotation, transcript and exon features of ENSEMBL were used to compute individual gene counts. Gene expression was estimated using RSEM (v1.2.30). Transcripts-per-million (TPM) was computed across samples and used in single-sample gene set enrichment analysis (ssGSEA) using R package GSVA (v2.0.7) and Hallmark gene sets from the Molecular Signatures Database (MSigDB).

### 2.2. Prompt generation

Figure 1 presents the prompt generation pipeline. We represented a histopathological whole-slide image by a set of spatially localized image tiles with *I*_*i*_ denoting the ith image tile extracted from a WSI, N is the total number of tiles. For each patient, bulk RNA sequencing represented by a vector of gene expression values G and the normalized enrichment scores for specific biological pathways derived from gene expression is represented by P:

**Figure 1.**
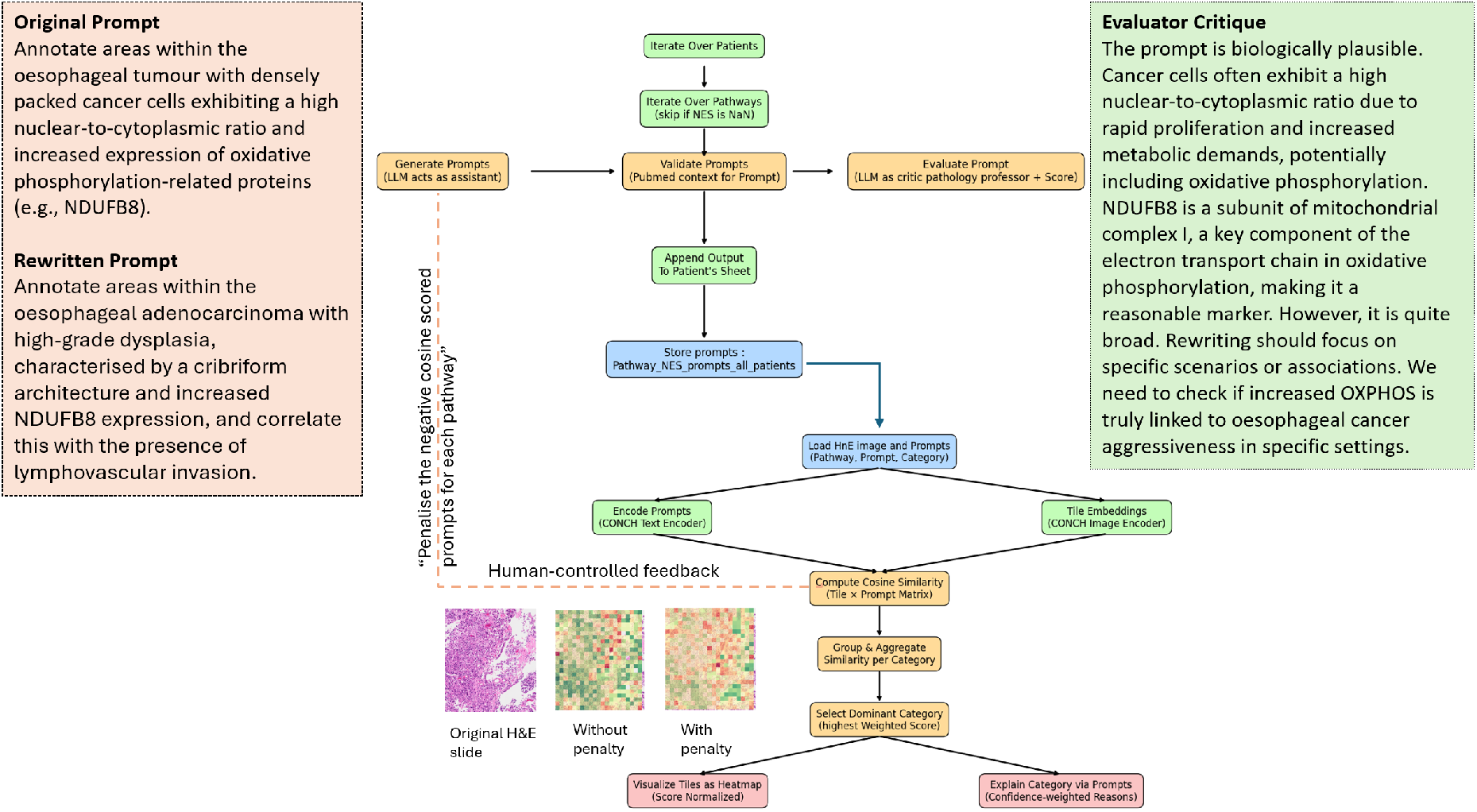
This schematic illustrates the human-in-the-loop CONCH pipeline for generating, evaluating, and visualizing biologically grounded prompts on histopathology slides. For each patient and each pathway with nonzero enrichment, approved prompts are stored and later loaded alongside the patient’s H&E image. The CONCH text encoder converts each prompt into an embedding, while the CONCH image encoder tiles the slide and produces corresponding image embeddings. Throughout, the Human may interject high-level guidelines to steer subsequent iterations.

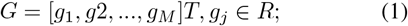

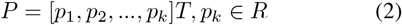

where, M is the number of measured genes and K is the total number of considered pathways and pathway *p*_*k*_ represents the integrated activity level derived from bulk RNA-seq data. Initial text prompts were generated using LLM model (L). Each prompt was conditioned on the structural prompt by: (i) assigning LLM as a pathology assistant (ii) the pathway of interest, (iii) its normalized enrichment score, and (iv) anatomical and oncological context specific to oesophageal cancer, (v) in subsequent iterations, a penalization mechanism was introduced whereby prompts from previous rounds were flagged and avoided during regeneration. Furthermore, each prompt generation cycle integrated human-authored feedback, captured from the user via terminal input, as part of a memory-guided prompting strategy. This structured feedback was prepended as instruction into the generative model to simulate expert intent refinement over multiple iterations. Mathematically, for image tile *I*_*i*_, to spatially localize these transcriptomic signals within histopathology images, we introduce generative large language models (L) parameterized as follows:

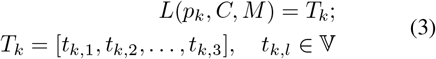

Here, *p*_*k*_ denotes pathways information, C represents the biological and morphological context represented as the patient specific system prompt, M indicates structured conversational memory with human feedback, and *T*_*k*_ is the set of generated textual prompts for the pathway *p*_*k*_ as shown in equation (3), with V denoting the natural language vocabulary space.

### 2.3. Evaluation of prompt-image alignment and critiquing

Each generated prompt was embedded using the pretrained foundation model, ensuring alignment in representational space with the pathology tiles. We used CONCH visual encoder *fv*(*·*) to embed histology tiles into a feature space and a pretrained textual encoder *ft*(*·*) to embed generated textual prompts into a compatible semantic space. Cosine similarity was computed between all prompt embeddings and tile embeddings. The semantic alignment between each tile embedding *e*_*i*_ and each prompt embedding *u*_*k,l*_ from both functions is quantified via cosine similarity:

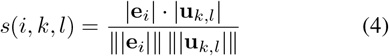

These similarity scores served as representations for morphological alignment: a higher score indicated stronger morphological resonance between the text-derived concept and the visual representation. To iteratively refine prompts, we incorporate a penalization function *P* (*t*_*k,l*_, *M*) based on historical conversational memory and human-in-the-loop feedback. Prompt evaluation also included PubMed-based semantic grounding: for each prompt, relevant biomedical abstracts were retrieved, and the same LLM model was assigned a different the role of pathology expert to critique and potentially revise each prompt written by pathology assistant LLM (*L*_*c*_) based on scientific evidence.

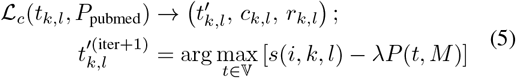

where *λ* is a hyperparameter controlling penalization strength using system-based prompts and assigned to penalize those prompts that have been assigned the negative cosine score. 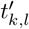 denotes the rewritten prompt based on biomedical evidence *P*_*pubmed*_,and *c*_*k,l*_ is its semantic category, and *r*_*k,l*_ ∈ [1, 5] indicates its quality score, which are categorized into three biological themes, proliferative instability, tumour suppression, and tumour aggressiveness, and scored on a 5-point scale for quality. Ultimately, the proposed optimization objective is to maximize the semantic alignment between image tiles and rewritten textual prompts over iterative refinement steps subject to domain knowledge-based constraints.

### 2.4. Visualization and Feedback-Driven Refinement

Tile-wise cosine similarity scores were aggregated to identify the dominant biological category per tile, computed as a weighted score combining average similarity and frequency of relevant prompts. Heatmaps were generated to visualize these dominant categories, highlighting spatially coherent regions where a given biological process may be active. By computing each tile’s score as the product of the number of categories-specific prompts and their mean cosine similarity, we simultaneously capture both the frequency with which a biological theme is invoked and the strength of its semantic alignment to tissue morphology. Mapping these weighted scores to a continuous color scale (e.g. green for low values through red for high) accentuates subtle spatial heterogeneity: green tiles mark regions with sparse or weakly supported hypotheses, while red tiles pinpoint robust hotspots of pathways activity. The iterative loop was executed up to a predefined maximum number of cycles. Terminating was optionally governed by visual convergence or prompt stability, as observed by the user. For each patient, radial plot included a summary of top-3 highconfidence prompts per tile and pathway, associated cosine similarity scores, and visual heatmaps.

### 2.5. Comparison of different LLMs and pathologist evaluations

The performance of different generative models (Gemini and Claude) is compared based on their cosine similarity scores and prompt diversity across patients and pathways. we conducted a qualitative assessment by comparing whole-slide image–based segmentations performed by a pathologist against those generated by our proposed model. For detailed evaluation of zoomed pathological regions, we implemented a quantitative scoring system: reviewer rated the tile-based heatmaps for tumor avidity (red = high avidity to green = low avidity) on a 1–5 scale, where (1) strongly uncorrelated, (2) uncorrelated, (3) neither correlated nor uncorrelated, (4) correlated, and (5) strongly correlated. Respondent were also invited to note any concerns or additional remark.

### 2.6. Quantitative evaluation

The model’s predictive performance was evaluated using the ablation study. First we run the training on a random subset of tiles per whole slide imaging versus training only on the identified hotspot tiles. We compare the results using the pearson correlation coefficient (PCC) between the pathway activities across. For each (Tile-Index, Pathway) pair, the PCC was computed between the vectors of cosine scores. The PCC measures the linear correlation between the two sets of scores. By directly comparing the PCC scores obtained from models trained on random tiles versus those trained on hotspot tiles, we quantitatively evaluated whether focusing on spatially localized, model-prioritized regions improves the accuracy and concordance of pathway activity predictions from whole-slide histopathology data.

### 2.7. Validation using Spatial Transcriptomic

We evaluated our approach on two datasets with ST ground truth: (i) an in-house oesophageal cancer patient with a 16×16 spot grid (Visium CytAssist, 10X Genomics), where adjacent spots were merged to match our 224×224 input resolution and corresponding expression profiles were averaged; and (ii) the public HEST prostate cancer patient [6], where spots/tiles are natively 224×224, requiring no averaging. For both, we (1) aggregated spot-level expression within spatial regions to construct a bulk sequencing input, (2) computed pathway activation scores from this bulk profile, (3) fed the bulk-derived pathway scores together with the spatial WSI tilescoordinates into our model, (4) extracted model-predicted tiles and validated them against the ST measurements, and (5) quantitatively compared prompts scores across ST regions using the quantitative metrices.

For each pathway, we represented every generated prompt as a sparse concept set derived from an explicitly fixed vocabulary (targets, morphology, cell types, processes, spatial cues, gene tokens, and a directionality). We embedded each prompt into a binary concept vector (one dimension per vocabulary term plus indicators for has gene and has directionality). We averaged vectors within ST and within bulk to obtain two centroids and calculated the centroid cosine, which summarises the overall semantic alignment between the two prompt sets at the pathway level.

In addition, the best-match Jaccard is computed by pairing each ST prompt with the single bulk prompt that maximises Jaccard overlap of concept sets.

## 3. Results

### 3.1. Integrated human-authored context

Figure 2 compares the role of integrating domain-specific human expertise in guiding large language models toward the generation of contextually relevant prompts for spatial pathway inferring analysis. Histopathological images (Figure 2a) of a male patient diagnosed with Barrett’s oesophagus located at the esophagogastric junction (Siewert II classification).

**Figure 2.**
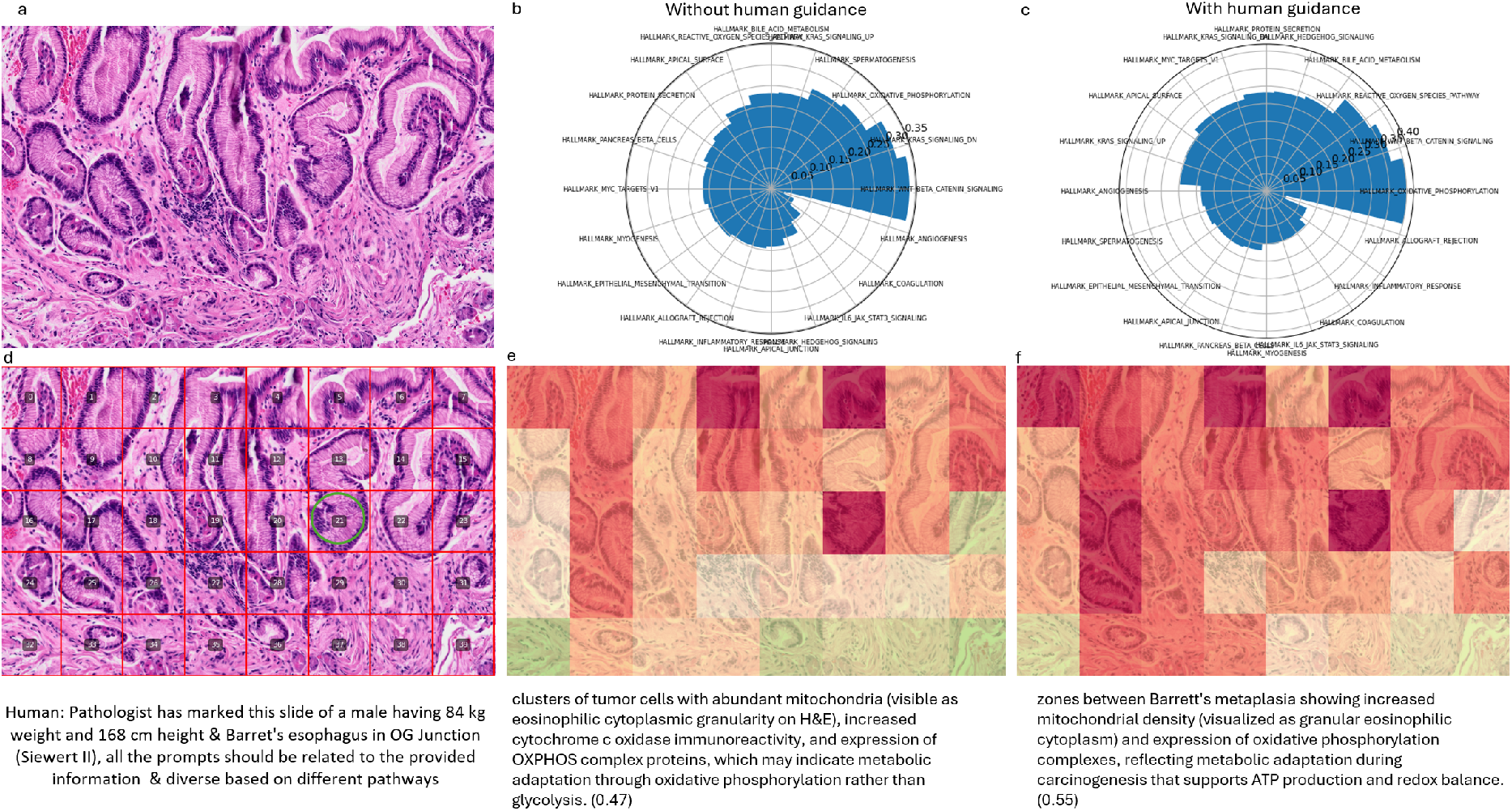
Illustration of the significance of human author-based domain knowledge information for guiding the LLM with generating specific prompts. A) Patient’s histopathology image. B) and C) Radial plots showing the comparison averaged cosine scores for the top three generated prompts without and with integrated human author context. D) shows the tiles overlayed for which the prompts are generated. E and F show the spatial heatmaps representing the avidity of the regions where biological hypotheses were weakly to strongly supported (from green to red) for without and with human guidance.

Radial plots (Figure 2b and 2c) illustrating a comparative analysis of averaged cosine similarity scores of the top three LLM-generated prompts, contrasting scenarios without and with integrated human-authored domain-specific context. The radial plots highlight substantial improvements in cosine similarity scores following human guidance, particularly in pathways critical for adenocarcinoma progression. As exemplified by tile 21 (Figure 2d green circled), two generated prompts are presented. Without human context (b), the generated prompt achieves an averaged cosine similarity score of 0.47, increased cytochrome c oxidase immunoreactivity, and expression of OXPHOS complex proteins, which may indicate metabolic adaptation through oxidative phosphorylation. In contrast, with integrated human-authored context (c), the prompt achieves a markedly higher score of 0.55. The enhanced prompt specifically highlights showing increased mitochondrial density (visualized as granular eosinophilic cytoplasm) and expression of oxidative phosphorylation complexes. Figures 2e and 2f show the spatial heatmaps representing the avidity of the regions where biological hypotheses were weakly to strongly supported (from green to red) for without and with human guidance.

### 3.2. Comparison between language models

Figure 3 presents a comparative analysis between Claude 3.7 and Claude 4.0 in generating biologically relevant prompts based on histopathology images across different hallmark pathways and patient tiles. The comparison is illustrated for poorly differentiated oesophageal cancer cases, each represented by histopathological tile maps and corresponding cosine similarity heatmaps for the generated prompts. Specifically, Claude 4.0 exhibits enhanced discriminatory capability and greater diversity in tile-wise spatial scoring. We have also compared other models for example, Gemini’s gemini-2.0-flash-thinking-exp-1219, gemini-learnlm-2.0 and gemini-2.5-flash-preview-04-17 and found gemini-2.5 average cosine score close to Claude 3.7 but flash Gemini 2.0 thinking model closer to Claude 3.7 for discrimination. In all other patients, Claude further substantiates its superior interpretative resolution, as indicated by clear demarcation of biologically active regions, whereas Gemini yields more generalized and lower-intensity heatmap patterns. Claude’s higher cosine scores suggest improved capability in capturing nuanced, pathway-specific morphological and molecular characteristics from histopathological imagery (refer to Supplementary Figure 1s and 2s).

**Figure 3.**
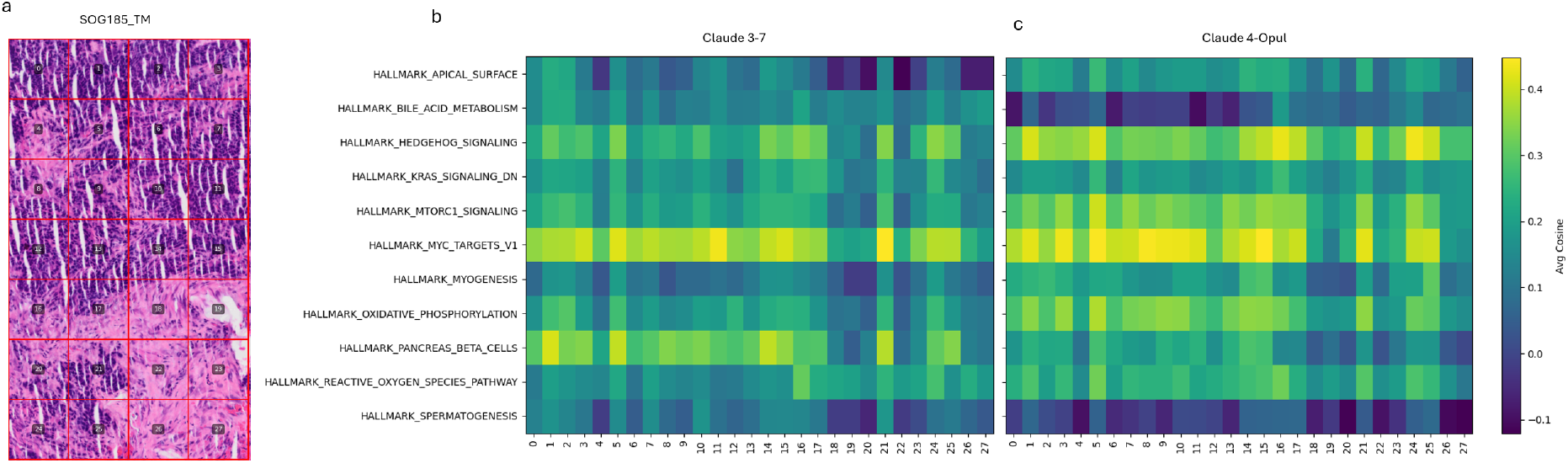
Comparison between Claude 3.7 and Claude 4.0 Sonnet model across the tiles and pathways for the average prompt’s cosine score. Claude 4.0 shows the higher score and diversity across the tiles scoring.

### 3.3. Comparison of spatial regions across pathology morphologies

Figure 4 illustrates heatmaps overlaid on H&E slides from oesophageal carcinoma samples, interpreted via hypotheses generated using the Claude 4.0 LLM. In the heatmaps, green tiles signify regions with sparse or weakly supported hypotheses, whereas red tiles highlight regions of pronounced pathway activity. Qualitative clustering observed from these spatial heatmaps demonstrates coherent regional consistency across histopathological subtypes. For example, Figure 4a demonstrate tissue from moderately differentiated oesophageal adenocarcinomas. It exhibits a patchy distribution, denoting heterogeneity in pathway activity with discrete islands of both heightened and reduced activity. Such spatial distributions reflect plausible variations in tumor biology at the microscopic scale, aligning with known complexities inherent in moderately differentiated adenocarcinomas, such as tumor nests showing increased mitochondrial content, evidenced by cytoplasmic eosinophilia, suggesting enhanced oxidative phosphorylation activity potentially related to PGC1*α*-mediated metabolic reprogramming due to oxidative phosphorylation pathway with a score of 0.67. Figure 4b portrays a poorly differentiated oesophageal adenocarcinoma with marked hotspots with intense red coloration are suggestive of aggressive tumour biology, prompted to show architectural disorganization, nuclear pleomorphism, and increased mitotic figures, consistent with MYC-driven cell cycle acceleration and proliferation (0.45). The clustered patterns were found to be similar to the human labelled patterns. Figure 4c corresponds to squamous mucosa from a case of squamous cell carcinoma. Notably, certain areas, predominantly localized toward the lower-left quadrant, manifest high activity hotspots. Here, the spatial heatmap distinctly prompted to show aberrant squamous differentiation patterns, particularly areas with nuclear accumulation of Gli1/Gli2 transcription factors or increased PTCH1 expression, which may indicate hedgehog pathway activation contributing to tumor progression (with 0.52 cosine similarity assigned for hedgehog signaling pathway in tile 10). Likewise, tile 21 in Figure 4d represents the Barrett’s metaplasia and adenocarcinoma at OG junction, characterizing areas with gradual cytological transition that show decreased KRAS pathway inhibitors (NF1, DAB2, RASA1) and regions with increased proliferative activity (Ki-67), indicating progression toward malignant transformation (0.55).

**Figure 4.**
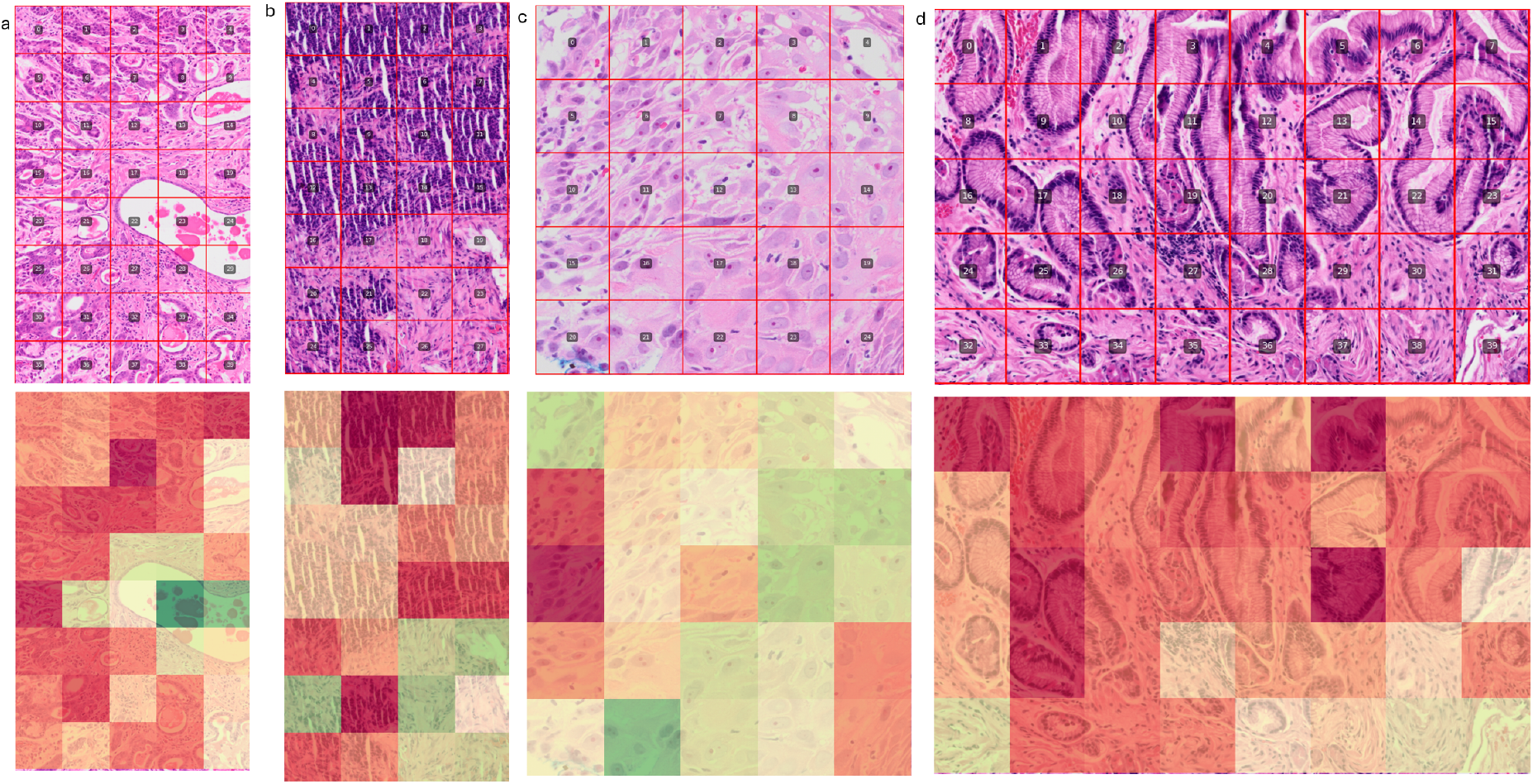
Representation of the spatial regions with sparse or weakly supported hypotheses, while red tiles pinpoint hotspots different of pathways activity using the Claude 4.0. Top row shows the original HnE slide with tiles overlay and bottom shows heatmaps. a) moderately differentiated oesophageal adenocarcinoma, b) poorly differentiated oesophageal adenocarcinoma, c) squamous mucosa for the squamous cell carcinoma and d) Barrett’s oesophagus. Green indicates regions where biological hypotheses were weakly supported. Red highlight areas of semantic alignment—potential hotspots of pathways activity.

Figure 5 shows distribution of prompt-cosine scores per pathway with low/high tile labels (as shown in previous Figure 4). This comparison of concentrated, high-scoring tiles against a pervasive low-scoring region underscores pronounced intratumoral heterogeneity and highlights the most informative areas for downstream spatial biology analyses. These comparisons illustrate that while CONCH reliably isolates both the most responsive and least responsive regions in each sample, the identity of these regions and the pathways they emphasize varies in a way that reflects the unique biology, grade and microenvironmental context of each tumor.

**Figure 5.**
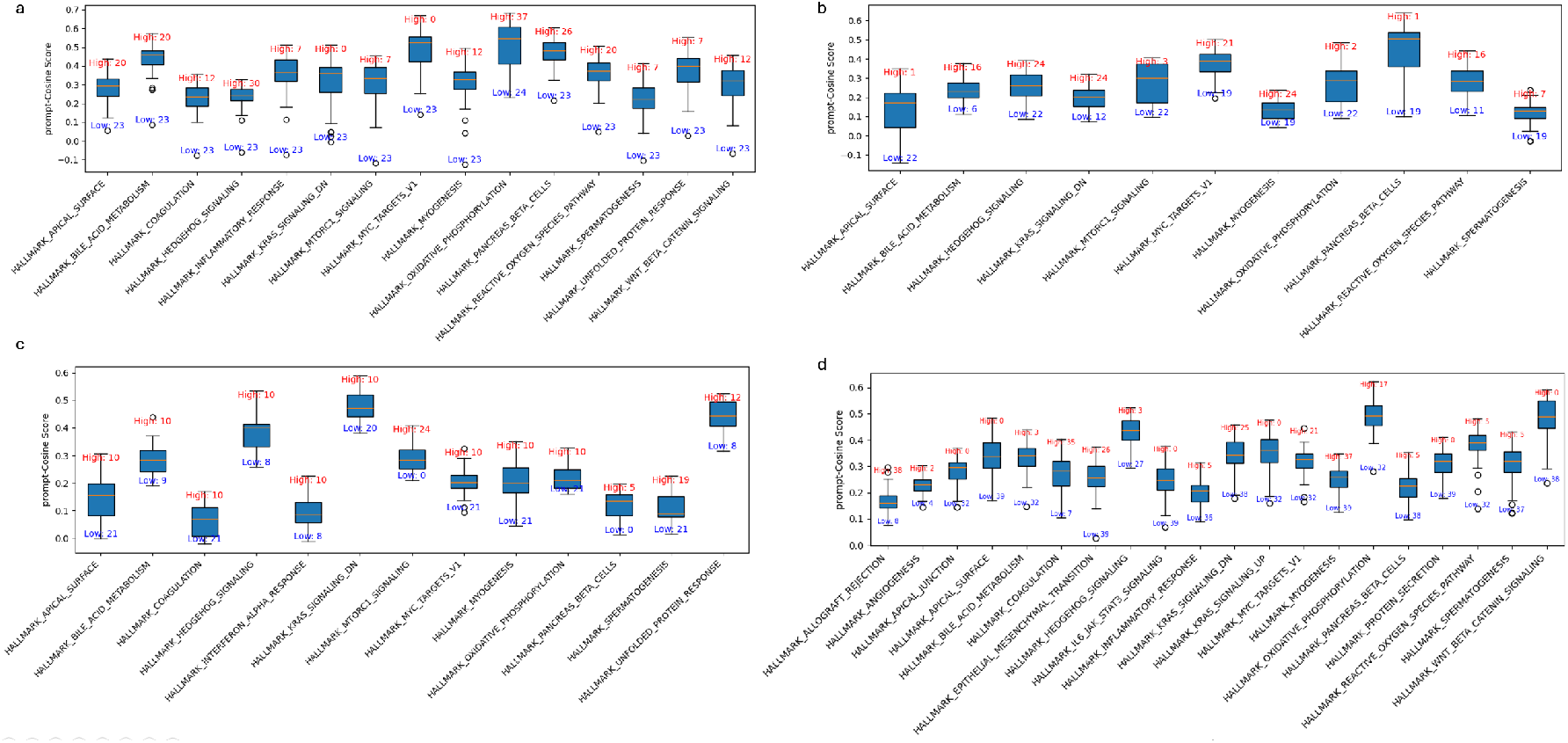
Distribution of Prompt–Cosine Scores per Pathway with Low/High Tile Labels (CONCH) for four whole-slide specimens. a) moderately differentiated oesophageal adenocarcinoma, b) poorly differentiated oesophageal adenocarcinoma, c) squamous mucosa for the squamous cell carcinoma and d) Barret’s oesophagus. For each hallmark pathway on the x-axis, the box-plot denotes the interquartile range of cosine similarity scores between image-tile embeddings and natural-language prompts. Red labels above each box identify the tile index with the highest score, while blue labels below denote the tile index with the lowest score.

### 3.4. Pathologist evaluation

Quantitative scoring was performed by the pathologist for detailed evaluation of zoomed pathological regions as listed in Figure 4. Across the reviewed cases, the tile-based heatmaps were rated highly (mean score 4/5), indicating overall concordance between color-coded avidity and underlying histology. These observations confirm that our model’s tile-based avidity heatmaps generally mirror the pathologist’s assessments. The sole moderate rating (scored 3 for Figure 4d) highlights areas for further refinement in capturing finer morphological heterogeneity.

Figure 6 Compares manual and automated segmentations across three esophageal pathology scenarios. Overall, the ensemble of heatmaps across these three pathological scenarios closely mirrors the pathologist’s manual annotations capturing both the location and relative intensity of disease regions. Squamous differentiation with intercellular bridges was clustered by the proposed method as identified by the pathologist with (Figure 6a) well defined cell borders. In the Barrett’s esophagus specimen (Figure 6b), the expert pathologist’s manual segmentation delineated two discrete glandular regions. Our proposed model closely replicated these boundaries. For the moderately differentiated adenocarcinoma case (Figure 6c), the pathologist demarcated invasive tumor nests. The model’s segmentation heatmap successfully captured these invasive fronts, assigning high-avidity scores to the tumor nests with minor under-segmentation occurred at the periphery of smaller nests. In the poorly differentiated adenocarcinoma specimen (Figure 6d), manual annotation by the pathologist encompassed a broad, irregular tumor mass with indistinct margins. The automated heatmap likewise highlighted this diffuse tumor region with intense red signals, accurately reflecting areas of high cellularity and metabolic activity. This strong visual and quantitative agreement underscores the proposed model’s potential to assist in rapid, reproducible tissue segmentation in diverse histopathological contexts.

**Figure 6.**
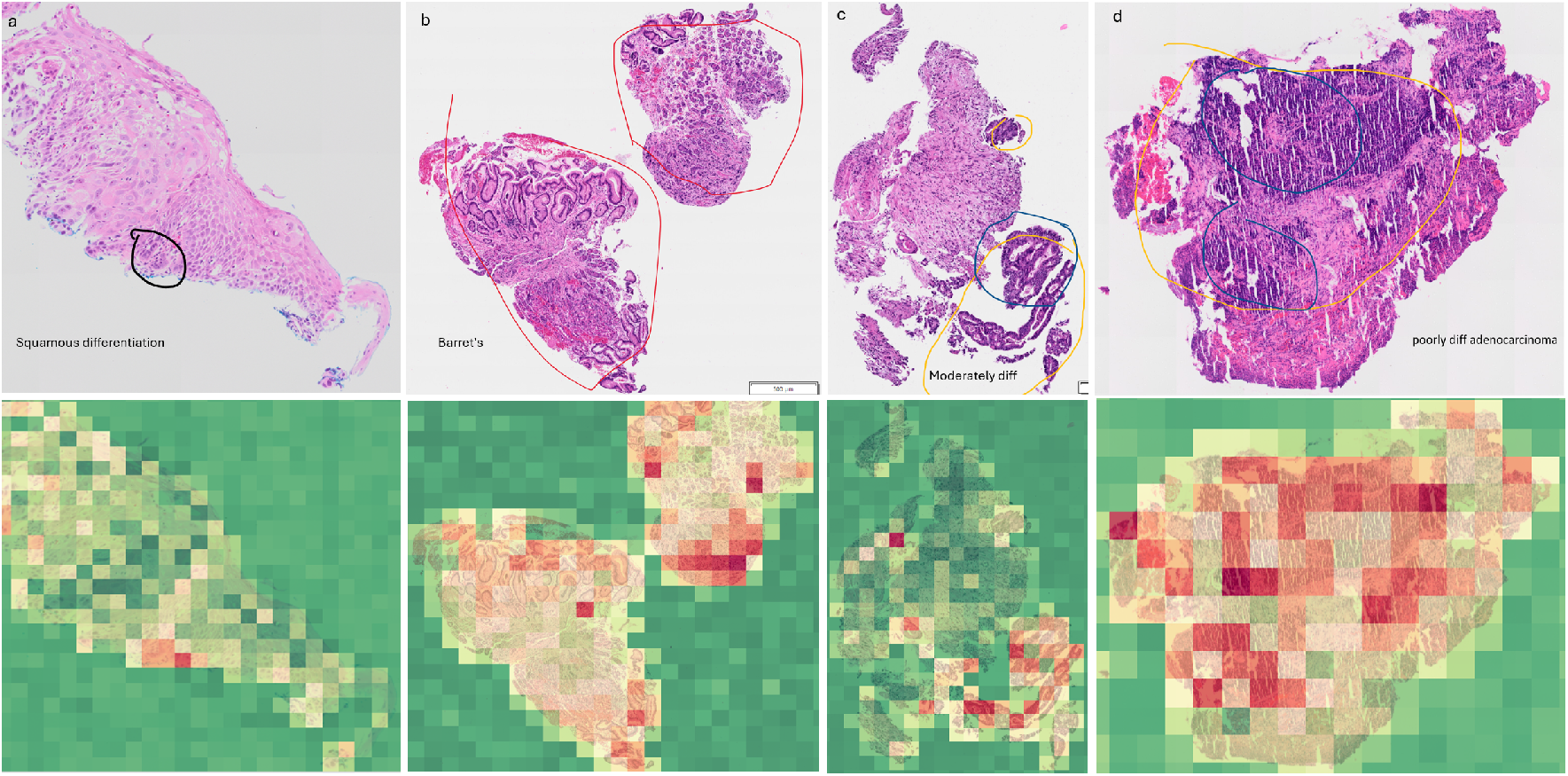
Comparison of manual and automated segmentations across four oesophageal pathology scenarios. Top row (a–d): whole-slide images with manual region annotations by an expert pathologist (outlines). (a) Squamous differentiation with intercellular bridges and well-defined cell borders, (b) Barrett’s oesophagus with two discrete glandular islands of outlined in red. (c) Moderately differentiated adenocarcinoma with tumor nests interspersed with cancer, highlighted in yellow. (d) Poorly differentiated adenocarcinoma with broad & irregular tumor mass with indistinct margins delineated in yellow and blue. Bottom row (a-d: Corresponding tile-based heatmaps produced by the proposed method. In each panel, the regions of elevated heatmap intensity spatially coincide with the pathologist’s manual annotations, demonstrating strong concordance between automated and expert segmentations in all three pathological contexts.

### 3.5. Quantitative Evaluation

Figure 7 presents the distribution of the pearson correlation coefficient of two ablation studies. For each patient, the PCC values remain consistently high (median values above 0.85), indicating a strong linear association between model predictions and ground truth across both sampling strategies. The narrow interquartile ranges of PCC further demonstrate the robustness of correlation for most pathways.

**Figure 7.**
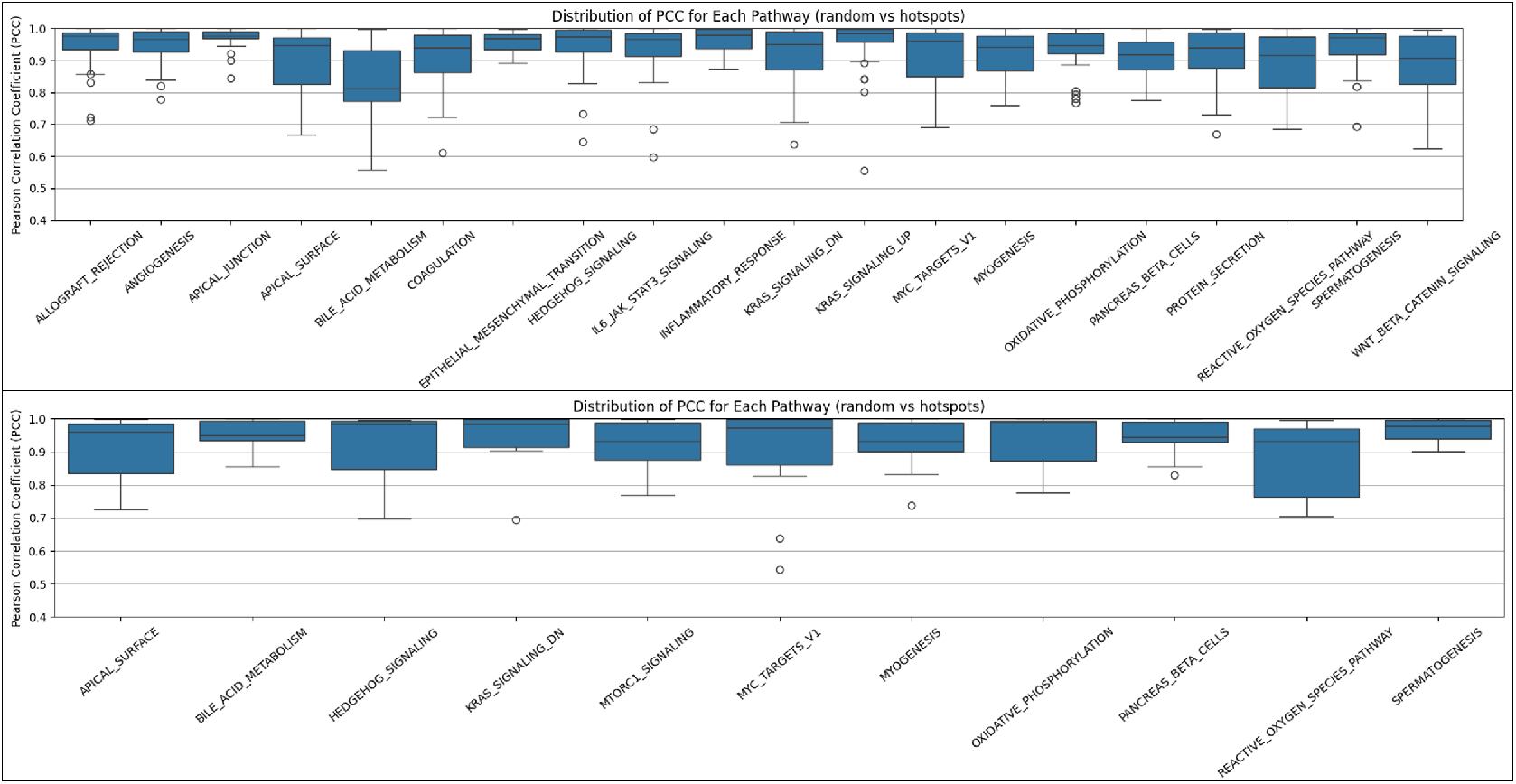
Distribution of the pearson correlation coefficient comparing model performance when trained on random tiles versus hotspot tiles identified by the proposed method for Barrett’s and poorly differentiated adenocarcinoma (as shown in figure 4d and 4b respectively)

Figure 8 demonstrates the average percentage distribution of tile classifications using the Claude LLM across 130 patient histopathology samples, classified into three biological categories: tumour aggressiveness, tumour instability, and tumour suppression. Oesophageal adenocarcinoma, squamous cell carcinoma and Barrett’s high-grade dysplasia exhibited the highest proportions of tiles classified as tumour aggressiveness, with substantial contributions also from instability-related features, indicating strong malignant potential. Necrosis, Barrett’s oesophagus and normal squamous mucosa demonstrated higher proportions of suppression-associated tiles, consistent with their less malignant phenotype, although instability remained present in varying degrees. Collectively, these findings highlight distinct molecular and pathological behaviour across oesophageal disease subtypes, with malignant states skewed toward aggressiveness and instability, whereas premalignant and non-malignant tissues retain greater representation of suppression.

**Figure 8.**
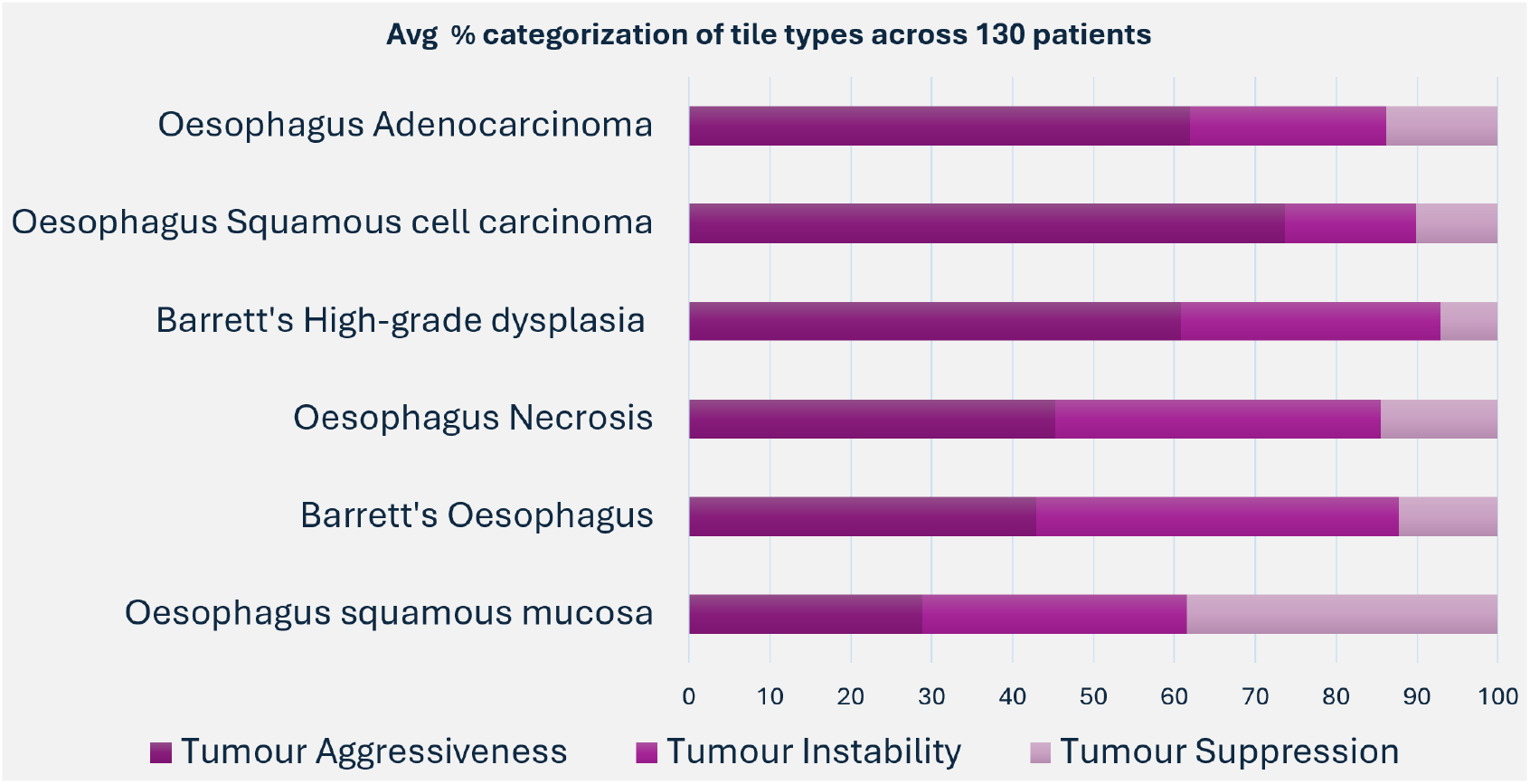
Average percentage classification of tile categorizations across 130 patients

### 3.6. Spatial Transcriptomic Validation

Figure 9 shows the randomly selected 9 spots on the WSI for prostate cancer HEST data. Qualitatively, our method produced red, yellow, and green-coloured annotations representing tumour aggressiveness, unstable, and suppressive regions (shown in blur). Figure 9b demonstrates clear spatial heterogeneity in pathway activation across the WSI. Box plots of the Hallway Pathway Score for tiles 0–9 show distinct distributions with different medians and dispersion, indicating that some regions capture bulk-like behaviour while others deviate substantially. Figure 9c shows that centroid cosine values are generally high (mean 0.7), implying that ST and bulk prompt sets, when averaged within each tile, emphasise broadly similar morphology. In contrast, best-match Jaccard scores are lower and more dispersed as shown in 9d, meaning individual ST prompts don’t duplicate bulk counterparts. Tiles with high cosine and elevated Jaccard contain highly morphology where both prompt sets converge. Tiles with high cosine but low Jaccard capture the same theme using diverse descriptors and tiles with low cosine and low Jaccard indicate semantic drift. Overall, trend shows that the average inference emphasized by ST prompts are closely aligned with those from Non-bulk prompts. A similar trend was observed in the tile 5 and tile 6 of OC cancer type (as shown in Supplementary Figure 5s)

**Figure 9.**
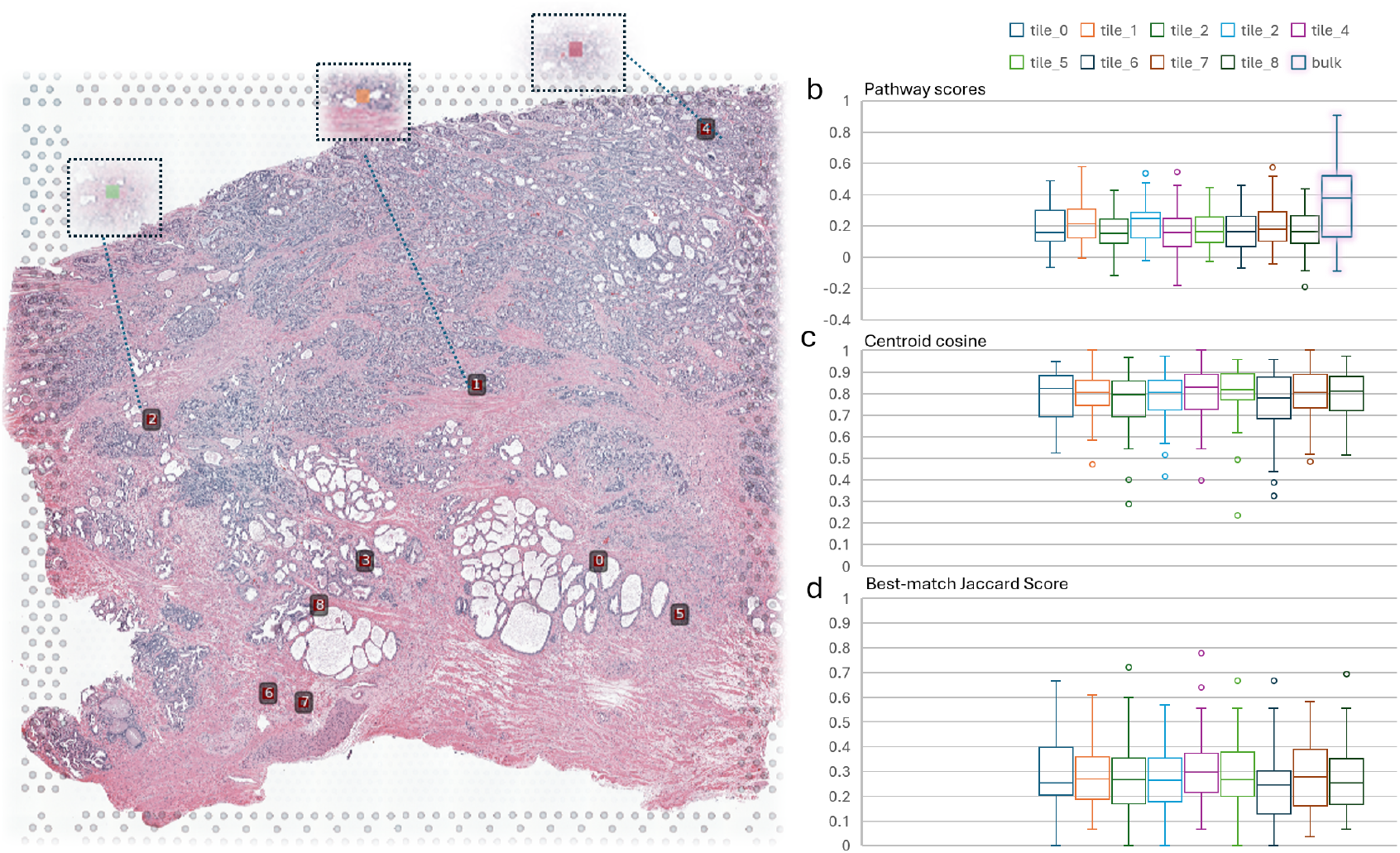
(a) ST Validation on WSI from the HEST prostate cohort with nine randomly selected tiles (0–8) highlighted; insets show qualitative annotations. (b) Tile-wise distributions of the Hallway Pathway Score compared with bulk. (c) Centroid-cosine between ST-conditioned and bulk-conditioned prompt sets per tile (higher = greater global semantic alignment). (d) Best-match Jaccard between individual ST prompts and their closest bulk prompts per tile

## 4. Discussions

We propose an AI agent that integrates a cyclic, human-in-the-loop workflow aimed at iteratively refining AI-generated prompts and critiques within a biomedical research context, specifically for applications involving bulk RNA sequencing based computational spatial pathway inferring and histopathology analyses. The proposed approach comprises literature retrieval, text summarization, critique evaluation, and memory-driven iterative refinement, thus establishing a comprehensive, self-improving system.

Unlike existing methodologies, our approach systematically records expert feedback and prior AI-generated outputs, enabling the AI agent to utilize comprehensive historical context during each iteration (refer Figure 1). This structured approach ensures that iterative refinements are context-aware and progressively enhanced. The integration of explicit summarization and critique stages within each cycle ensures each refinement step is carefully validated and informed by expert knowledge, thereby improving biological nuance and spatial accuracy. Figure 2 underscore that the use of targeted, pathology-specific human insights substantially refines the biological relevance of LLM outputs. Expert-guided prompting strategies can significantly improve the interpretability and clinical applicability of AI-driven spatial analyses. For example, in Figure 4 pathologist noted minimal visual distinction among neighbouring tiles—particularly in the bottom-right (tiles 19 vs. 13) and top-right (tiles 10 vs. 6) quadrants—indicating that for ambiguous morphology, the heatmap may require further expert-guided prompting to enhance sensitivity.

In Figure 5, the prompt–cosine score distributions share a common pattern, each pathway yields a characteristic range of alignment values, and extreme cold-spot (green) and hot-spot tiles (red) can be readily identified in Figure 4, yet the specifics of which biological programs dominate, and which regions underperform differ markedly between patients. In every case, a set of tiles acts as a pervasive coldspot, scoring lowest across most or all pathways. This suggests a non-neoplastic region that universally fails to match any molecular prompt. The consistency of this phenomenon confirms that each slide contains region relatively devoid of disease-specific morphological features.

We have tested the enhanced system prompt designed to convey granular, tile-level context. For instance, when evaluating the same MYC pathway score, the baseline prompt yields: “Infiltrative nests of poorly differentiated adenocarcinoma cells showing increased chromatin texture heterogeneity, multiple small nucleoli per nucleus, and peritumoral lymphocytic infiltrate depleted of *CD*8^+^ T cells characteristic of MYC-mediated immune evasion.” By contrast, the advanced prompt produces a more detailed description: “Superficial mucosal ulceration overlies sheets of discohesive adenocarcinoma cells with prominent nucleoli, abundant mitotic figures, and apoptotic debris. These tumor cells exhibit nuclear MYC overexpression accompanied by upregulated ribosomal protein expression, PKM2-driven glycolytic reprogramming, and elevated ODC1 levels, as they invade through a fragmented basement membrane alongside TGF-*β*-secreting cancer-associated fibroblasts.” In addition, Figure 3 compares the latest LLM by the anthropic (Claude 4-Opal) predicting comparatively distinct clusters in comparison to the Claude 3-7. Combinatorial effect of the designed system prompts and the LLM model used, affects the generated prompts, however the generated spatial heatmaps were found to be consistent.

Bulk RNA-seq combined with WSI approximates ground truth ST when the underlying biology is diffuse and as shown in Figure 9. In such cases, tile-level pathway scores closely overlap the bulk distribution and centroid-cosine values between ST-style and bulk prompts are high (0.85–1.0), indicating thematic agreement. In addition, best-match Jaccard remains modest.

Under these conditions, ST may be deferred for slide-level conclusions or coarse region selection by using Agent SPI-WSI on bulk+WSI as a cost-effective surrogate. It implies the proposed method can be used to screen slides, saving the overall costs. ST can be run only on slides/spots flagged as heterogeneous. Low centroid-cosine together with low best-match Jaccard identifies tiles whose semantics diverge from bulk, reflecting micro-niche biology (e.g., invasive fronts, perivascular cuffs, tertiary lymphoid structures) or compartmental mixtures that bulk cannot resolve.

Our approach integrates bulk RNA-seq–derived pathway scores with whole-slide images and uses prompt semantics to localize pathway activity at the tile level, with validation against spatial transcriptomics. In contrast to previous studies, GHIST and spot-level baselines (ST-Net, Hist2ST, His-ToGene, DeepPT, DeepSpaCE, GeneCodeR), which perform per-gene regression and typically require paired ST for training, our method does not predict gene-level expression. Consequently, per-gene metrics such as PCC/SSIM are not directly applicable.

Nonetheless, when Pearson correlation coefficient distributions were indirectly compared, our pathway-level results (Supplementary; median =0.20; IQR = 0.08–0.35; range = 0.01–0.40) occupy the same numerical band reported for gene-level ST prediction by state-of-the-art models (e.g., ST-Net, Hist2ST, DeepPT, DeepSpaCE, GeneCodeR, THItoGene, iStar, GHIST), whose violins typically centre around 0.10–0.20 with upper tails near 0.30–0.40 [3]. Although this is not an ideal comparison our estimates are pathway correlations obtained with bulk+WSI and no paired ST at inference, whereas the literature plots gene correlations learned with paired ST the overlapping ranges suggest that a bulk-anchored, prompt-based approach can achieve correlation behaviour comparable to spot-supervised methods while requiring markedly less supervision. We therefore interpret the present analysis as preliminary evidence of competitive performance.

In addition, the centroid-cosine between ST-conditioned and bulk-conditioned prompt sets is high for most tiles (median in the high-0.7 range), indicating that in a morphology-aware embedding space the average semantics of the two prompt families are closely aligned. This again mirrors GHIST’s observation that histology–transcriptome agreement is strongest for spatially informative signals and weaker for globally diffuse programs [3]. By design, our centroid-cosine functions as a morphology-weighted concordance, capturing alignment where tissue architecture is stereotyped. In contrast, the best-match Jaccard is lower and more variable, reflecting micro-environmental heterogeneity and instance-level variability: individual ST prompts seldom have near-duplicates among bulk prompts even when their centroids agree. Taken together, these results (Figure 9a) indicate that bulk+WSI in-context prompting recovers much of the global ST signal while preserving local diversity; explicit spatial supervision remains valuable for fine-grained, tile-specific biology that bulk anchoring may under-specify.

There remains notable room for methodological refinement. First, the magnification of tile and shape delineation can impose constraints on biological interpretation, potentially obscuring subtle transitions in cellular states or microenvironmental factors. Future iterations may benefit from refined spatial granularity or continuous heatmap representations to enhance biological relevance. Second, domain expert based human feedback to guide iteratively have the potential to enable problem specific prompt generation for specific insights. In future work, we will be exclusively testing this robustness of the method. The selection of the tile size is one of the critical factors in the proposed work. We have utilised the pre-trained CONCH that was trained with the size of (224,224), reducing the tile size below this threshold does not account for the better discriminability. Eventually, other vision-language model in pathology can also be compared alongside CONCH. Additionally, validation against immunohistochemical or genomic benchmarks would substantiate and reinforce the interpretative accuracy of these qualitative clustering observations.

## 5. Conclusion

Overall, this approach exemplifies a robust methodology combining literature retrieval, targeted summarization, expert critique, and iterative conversational memory-based refinement, thus demonstrating significant potential for applications requiring nuanced biomedical interpretation and computational spatial correlation between gene expression data and histopathological observations. Agent SPI-WSI achieves bulk-equivalent pathway characterisation across the slide, as evidenced by overlapping score distributions and high centroid cosines. Our future work involves utilising reinforcement learning and chains of thought to dynamically incorporate micro-anatomical anchors and measurable morphology into system prompts.

## Acknowledgments

The authors wish to thank the patients who participated in our project. We acknowledge the work and support of the Upper GI Unit at Princess Alexandra Hospital, Brisbane, Australia. This research was partially carried out at the Translational Research Institute, Woolloongabba, QLD 4102, Australia. The Translational Research Institute is supported by a grant from the Australian Government. This study was funded by a large Cancer Council Queensland (CCQ) grant, aimed at Accelerating Collaborative Cancer Research. Lauren Aoude is supported by a National Health and Medical Research Council (NHMRC) Emerging Leadership 2 (EL2) grant (APP2034399). Rajat Vashistha acknowledges his fellowship from CCQ ACCR-190.

## Declaration of generative AI and AI-assisted technologies in the writing process

During the preparation of this work the author(s) used chatgpt 4.0 in order to improve the grammatical correction and readability. After using this tool/service, the author(s) reviewed and edited the content as needed and take(s) full responsibility for the content of the publication

## Appendix A

### Extended Methods

To systematically evaluate histopathological features linked to pathway-level transcriptional states, we developed an automated pipeline that integrates transcriptomic signatures, literature synthesis, and generative large language models to derive and critique prompts on a per-patient basis. The pipeline accepts normalized enrichment scores (NES) for multiple hallmark pathways across a cohort of patients,

Using the NES value for a given pathway, we query a generative large language model to synthesize concise, one-line pathology prompts. These prompts are designed to instruct vision-language models (e.g., CONCH) to spatially localize histological regions on H&E-stained biopsy slides that may reflect downstream consequences of the pathway’s dysregulation. The prompt generation is context-aware as shown in figure below:

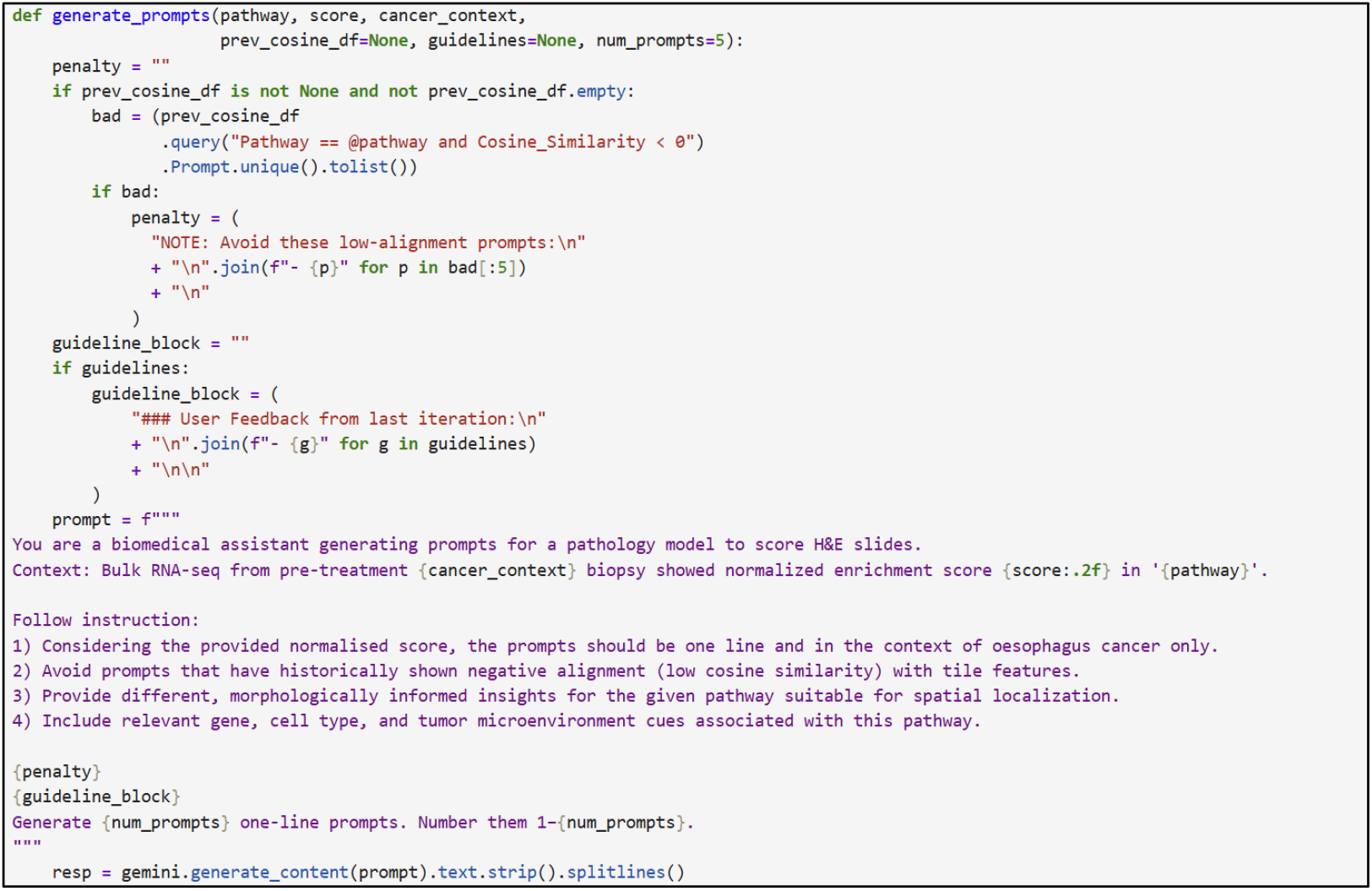

To assess the biological plausibility and histopathological relevance of each generated prompt, we perform a literature-guided critique. For each prompt, we construct a PubMed query and retrieve the top ten most relevant abstracts related to the pathway and cancer type. These abstracts are concatenated and provided as grounding context to a second call to the Gemini model, which is tasked with evaluating each prompt along four axes: scientific clarity, morphological interpretability, relevance to known TME-cell type-pathway associations, and suitability for spatial localization tasks. Prompts are either accepted or rewritten by the critic model, categorized into biological roles (e.g., tumor aggressiveness, suppression, or proliferative instability), and assigned a quality score as shown in figure below.

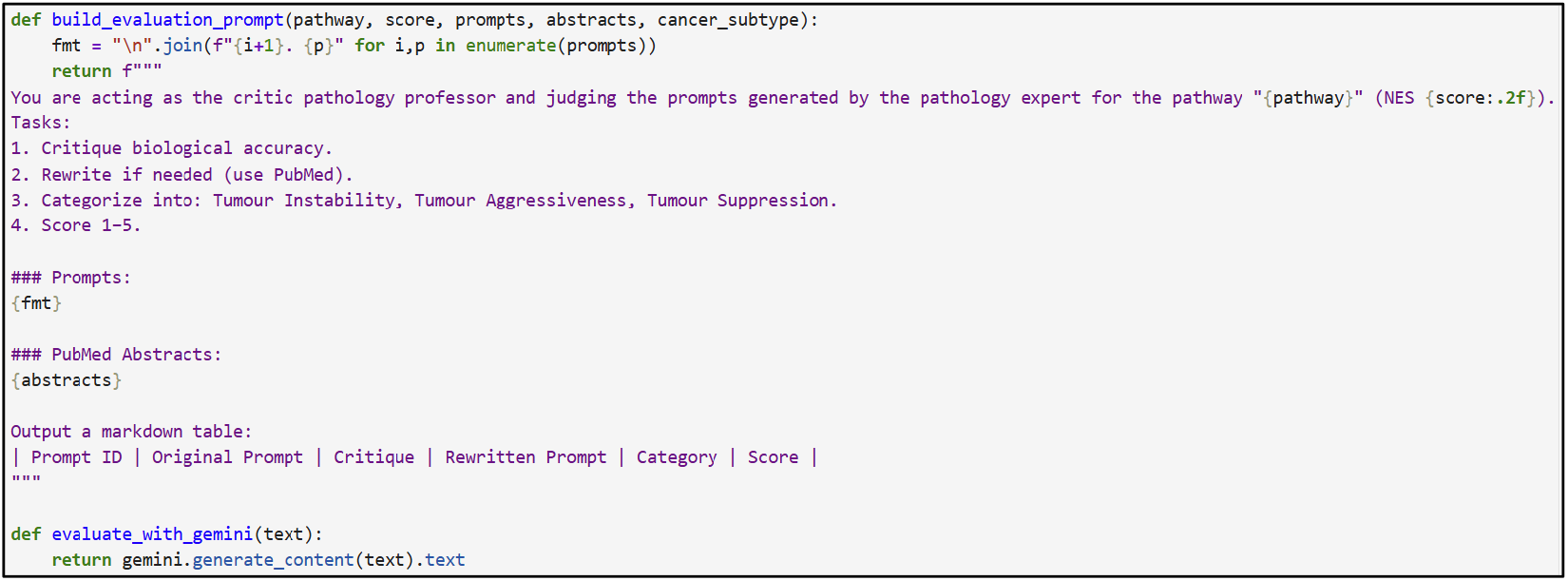

This multi-stage workflow enables biologically interpretable, pathway-informed semantic annotation of histopathology images at the tile level and supports downstream integration with spatial transcriptomics, clinical outcomes, or model training pipelines.

## Supplementary Figure

**Figure 1s.**
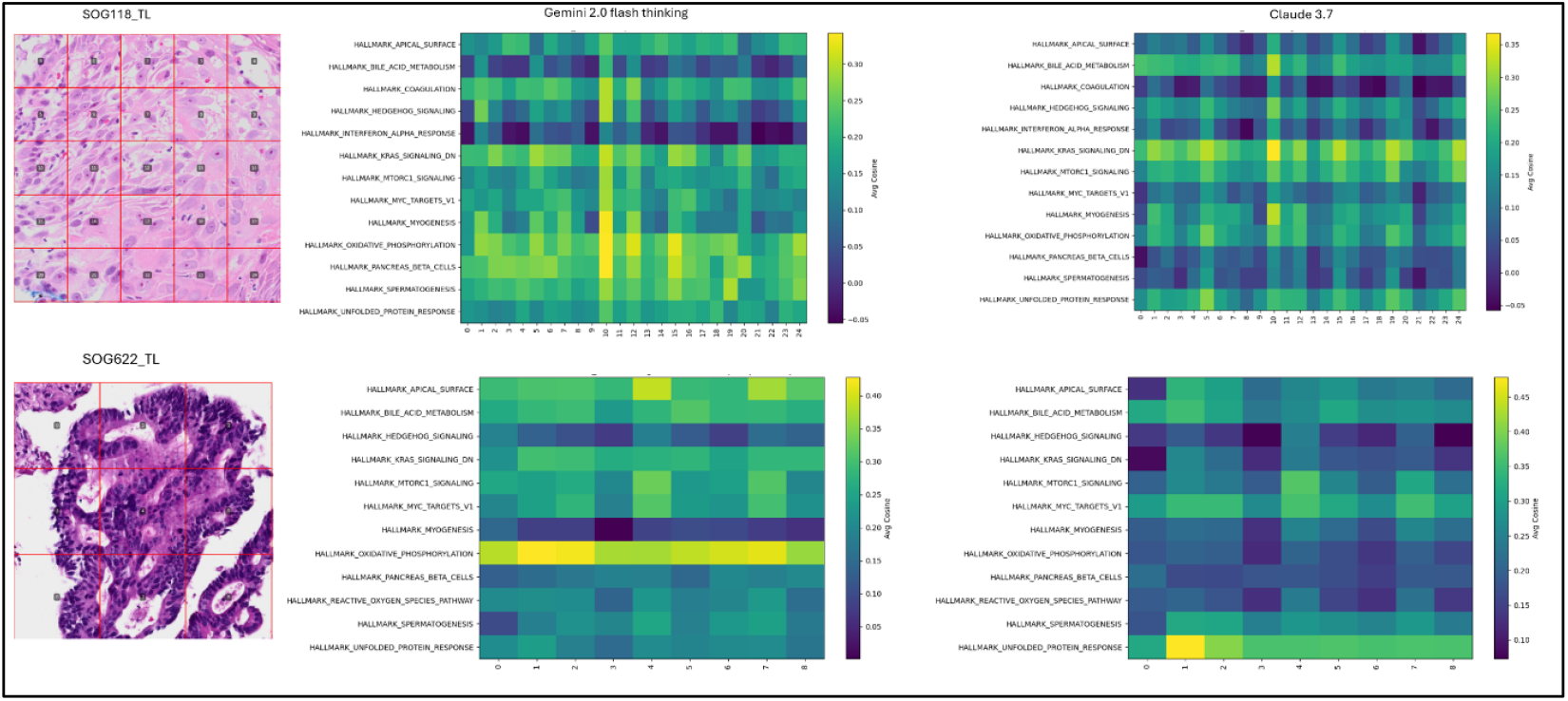
Comparative analysis between Gemini 2.0 Flash and Claude 3.7 Sonnet in generating biologically relevant prompts based on histopathology images across different hallmark pathways and patient tiles for two different patients

**Figure 2s.**
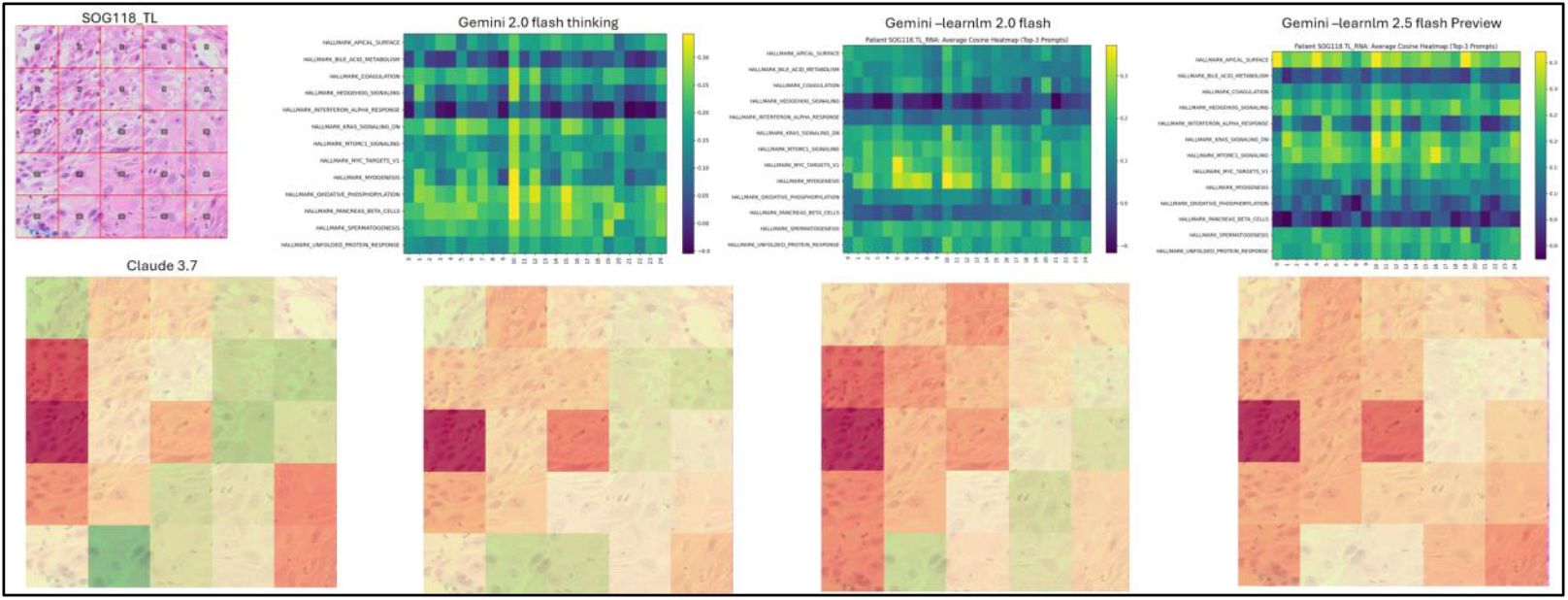
Comparison between Gemini’s gemini-2.0-flash-thinking-exp-1219, gemini-learnlm-2.0 and gemini-2.5-flash-preview-04-17.

**Figure 3s.**
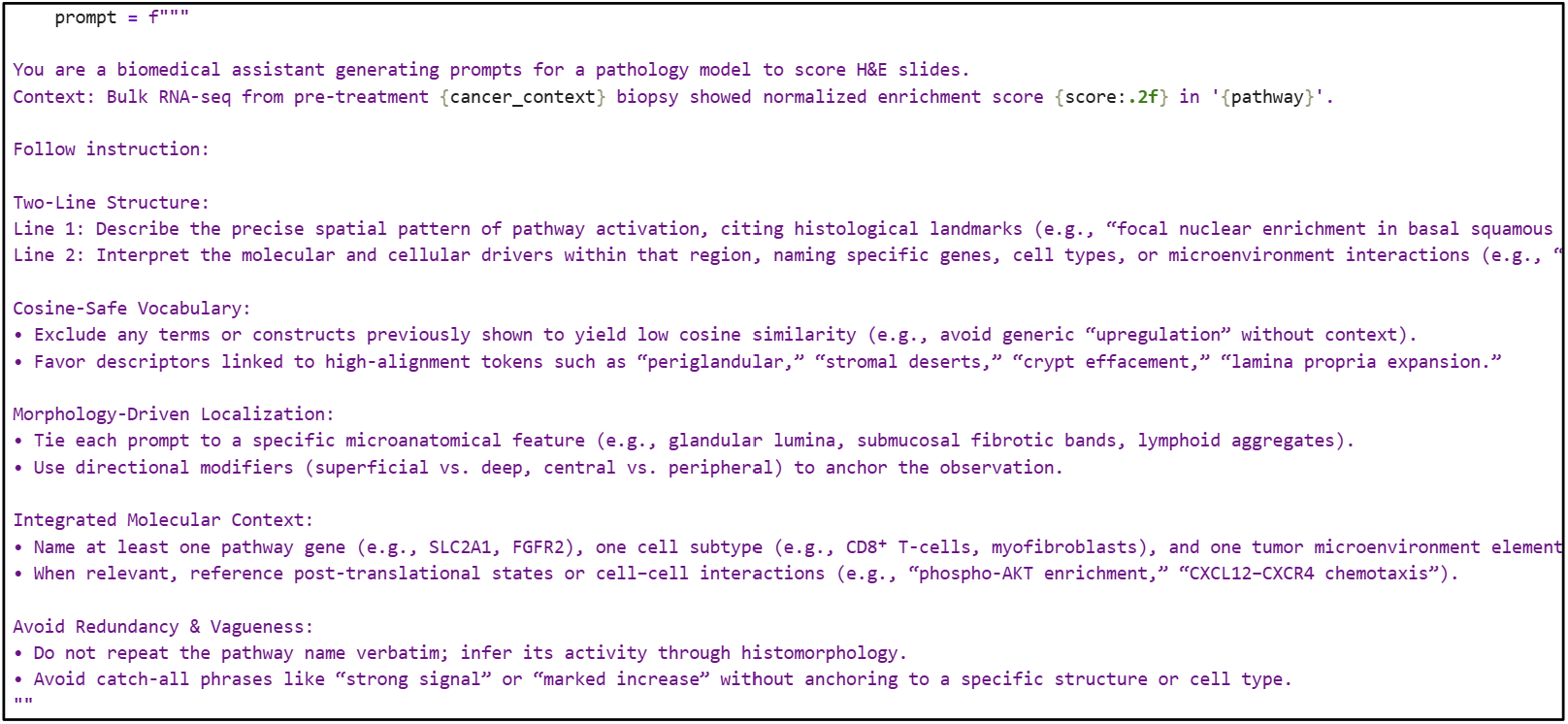
Enhanced system prompt

**Figure 4s.**
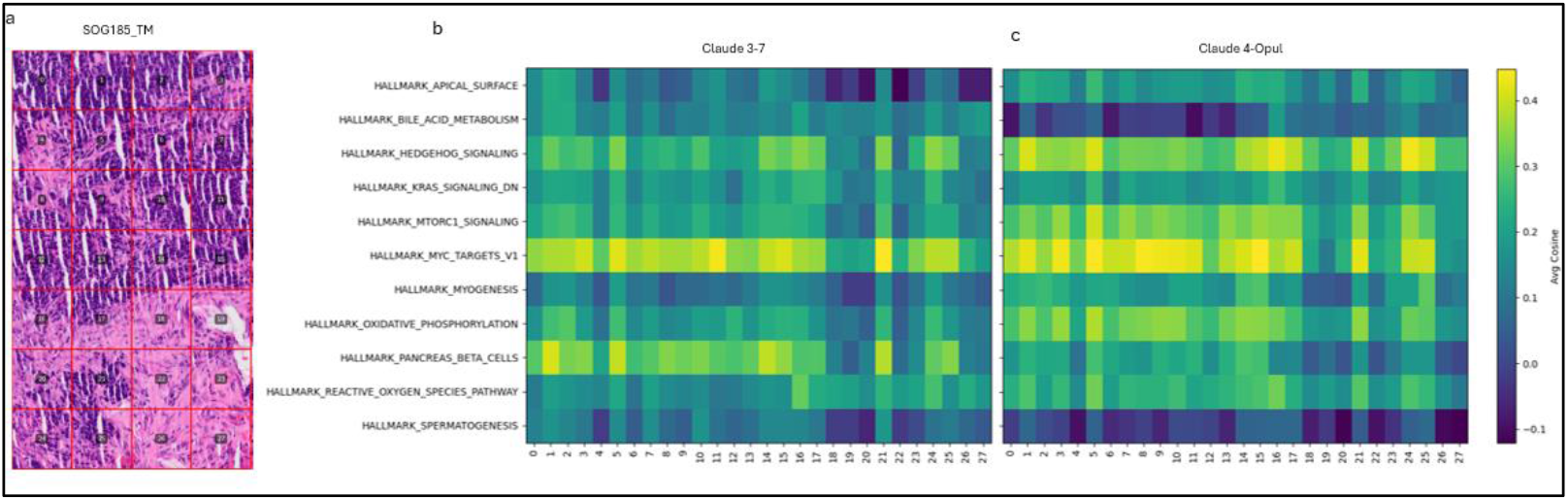
Comparison between version 3-7 and 4-opal by Anthropic.

**Figure 5s.**
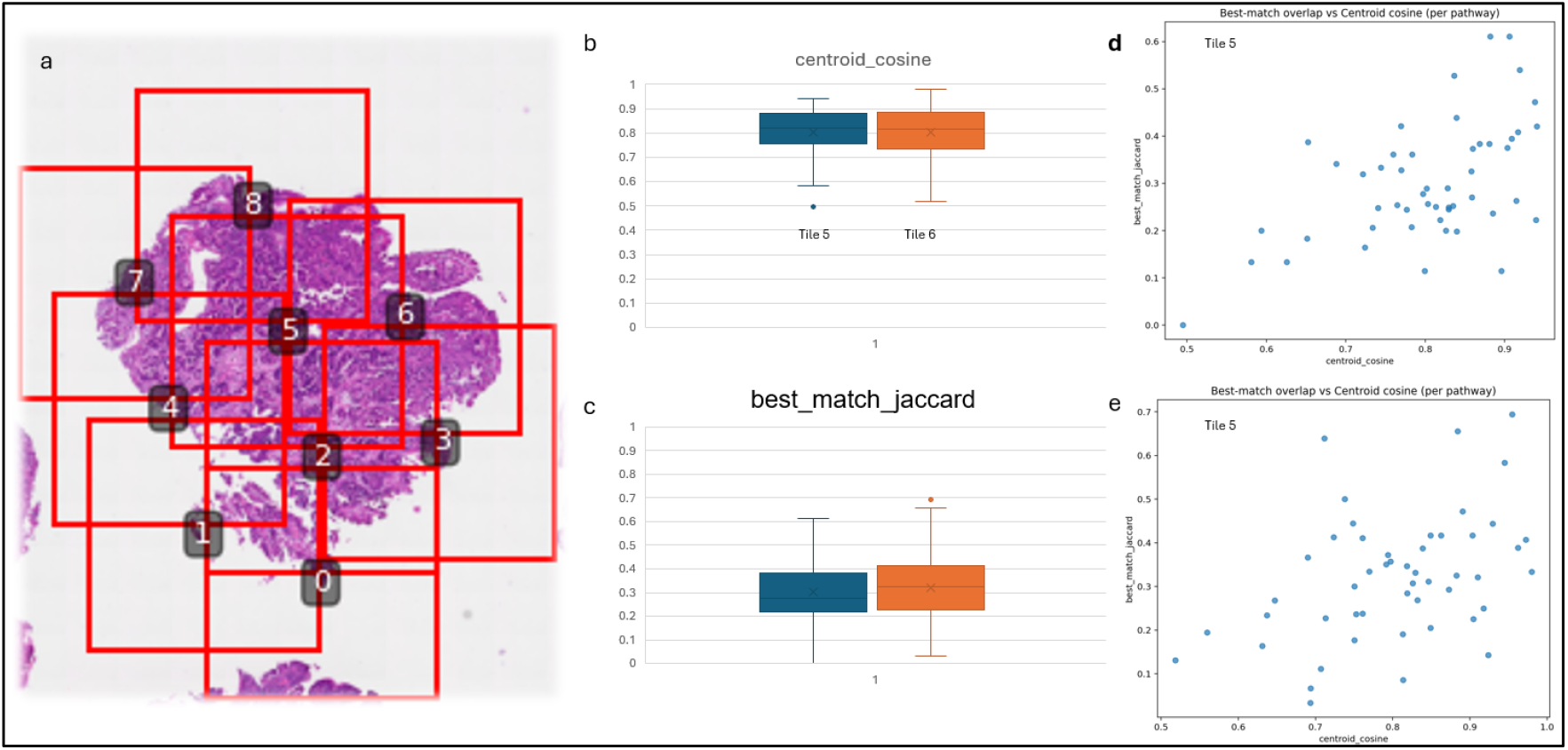
Whole-slide image (WSI) from the OC cohort with nine randomly selected tiles

